# Rare Variant Burden in Known Dystonia Genes in Population Controls and Sporadic Dystonia Patients

**DOI:** 10.1101/194399

**Authors:** Elizabeth T. Cirulli, Patrick Hickey, Samantha Tracy, Julia Johnson, Zachary Caffall, Kaylin Tsukayama, Burton Scott, Mark Stacy, Nicole Calakos

## Abstract

**Background:** Rare mutations in genes associated with Mendelian forms of disease are a potential mechanism for sporadic disease. The need to assess the clinical significance of such variants is increasing as personalized medicine and genome sequencing increases.

**Objective:** To evaluate the rate of rare, functional variants in dystonia genes in the general population to improve interpretation of the clinical relevance of potentially pathogenic variants in dystonia cases.

**Methods:** We performed an “aggregated” collapsing analysis of exome sequence that considered rare coding variants in genes previously associated with dystonia, a rare neurological movement disorder, on 2,372 population controls of European ethnicity. We then performed a pilot study in sporadic dystonia to assess whether there was a substantially greater incidence of individuals with rare variation in dystonia genes.

**Results:** Nearly half of population controls had a rare coding variant when 148 genes associated with a dystonia phenotype were considered. When the subset of genes causing isolated dystonia (14 genes) was evaluated, 3-4% of controls harbored rare qualifying variants. Our pilot study of case exomes was powered to identify a five-fold higher or greater rate of qualifying variants in isolated dystonia genes in sporadic dystonia cases compared to population controls; we did not find such an enrichment.

**Conclusions:** We provide the first systematic analysis of rare variation in dystonia genes considered collectively. Our findings emphasize the need to consider the overall frequency of variants in rare disease-related genes in the general population when considering their potential role in clinical presentations.

## INTRODUCTION

Human geneticists are increasingly appreciative of the role of rare variants in disease [1-4]. A role for novel mutations in genes associated with familial, Mendelian forms of diseases has been identified as a cause for sporadic presentations in a number of diseases, such as *de novo* mutations in Mendelian mental retardation genes [5]. Moreover, exome sequencing studies of sporadic cases in diseases such as amyotrophic lateral sclerosis and myocardial infarction have found support for contributions by both novel risk genes and known familial disease genes [6, 7], indicating heterogeneous genetic contributions.

Dystonia is an involuntary, centrally driven movement disorder involving slow twisting movements and sustained abnormal postures [8]. Dystonia can present alongside other neurological symptoms, as in the setting of neurodegenerative disease, metabolic disorders, trauma or stroke, or occur in isolation, as in the setting of drug side effects, rare inherited diseases, and the most common of isolated presentations, sporadic disease. Although a genetic component is clearly demonstrated in rarer forms of dystonia [9], much less is known about the etiology of adult-onset, sporadic disease, which includes presentations such as blepharospasm, cervical dystonia, and spasmodic dysphonia. Environmental, epigenetic, and genetic factors are all potential contributors.

In dystonia, rare coding variants in known dystonia genes have been reported in isolated sporadic cases [10-20]. In such isolated cases, a clinician may be tempted to attribute the rare variant to the patient’s clinical presentation. However, this approach suffers from ascertainment bias, since the individual’s frequency is “100%” *a priori*. Secondly, while large population databases are available to look up frequencies of rare variants, typically, either only the incidence of that specific variant or the overall incidence of variants in that specific gene are considered. As the finding of a rare missense variant in any gene associated with the patient’s clinical phenotype would have prompted making an attribution to their presentation, it is important to first consider the overall frequency of individuals in the general population that have similarly qualifying missense variants in disease-associated genes *considered collectively*. For example, to date, when the burden of rare variants has been considered, it has typically been assessed in single specific dystonia genes [11, 19, 21-23]. Thus, the overall likelihood of finding a rare variant in any dystonia-associated gene in the general population is not readily accessible to clinicians as yet because this has not been systematically evaluated.

In this study, we performed exome sequencing in 2,372 population controls and a small pilot cohort of 20 sporadic dystonia cases to determine the overall incidence of individuals with qualifying variants in dystonia genes. In this analysis, the presence of 1 or more rare missense variants in any dystonia-associated gene qualified an individual to be counted. Thus, rather than a gene-by-gene assessment, this analysis assesses the overall contribution of dystonia gene variation to population controls and sporadic cases. Importantly, this analysis does not consider the specific role of any given variant or gene in its association with dystonia cases.

This approach involves “collapsing” all variants in a group of genes for the purpose of assessing the overall population frequency of dystonia gene rare variation (reviewed in [24, 25]). This analysis also enables an assessment of whether rare variation in dystonia-associated genes occurs more commonly in cases than controls. Here, we provide a first assessment of this possibility using a small pilot cohort of twenty sporadic cases evaluated in our Movement Disorders specialty clinic.

## METHODS

### Samples

The study was approved by the Duke University Medical Center IRB, and all subjects gave written informed consent. Control samples were sequenced as part of other studies at Duke University Medical Center and were not enriched for (but not specifically screened for) dystonia or other neurological disorders (Table 2). Cases were recruited at the Movement Disorders Center at Duke University Medical Center, Durham, NC. We collected 20 unrelated patients diagnosed with adult onset, sporadic dystonia (3 men and 17 women) with a mean age of dystonia onset of 47.4 +/- 9.59 years. A diagnostic workup was conducted by a movement disorders specialist to confirm the symptoms of dystonia with muscle involvement classified as focal, segmental, multifocal, or generalized. Only presumptive primary cases were recruited. As is typical for our sporadic dystonia patients, no prior genetic testing had been performed for the majority (1 case had negative Dyt1 testing). Secondary dystonias associated with conditions such as Parkinson’s disease or other neurodegenerative disease were excluded. Cases suggestive of Mendelian inheritance were also excluded. A complete family and medical history was collected including common toxic exposures and medical comorbidities.

### DNA sequencing

Genomic DNA samples from 2,372 population controls and twenty subjects with dystonia were exome sequenced at Duke University. Control samples were either exome sequenced using the Agilent All Exon (37MB, 50MB or 65MB) or the Nimblegen SeqCap EZ V2.0 or 3.0 Exome Enrichment kit or whole-genome sequenced, and case samples were sequenced using the Agilent All Exon 37MB or 50MB kit, using Illumina GAIIx or HiSeq 2000 or 2500 sequencers according to standard protocols. All samples were processed using the same methods, as follows. The Illumina lane-level fastq files were aligned to the Human Reference Genome (NCBI Build 37) using the Burrows-Wheeler Alignment Tool (BWA)[26]. We then used Picard software (http://picard.sourceforge.net) to remove duplicate reads and process these lane-level SAM files, resulting in a sample-level BAM file that is used for variant calling. We used GATK to recalibrate base quality scores, realign around indels, and call variants [27]. Variants were required to have a quality score (QUAL) of at least 20, a genotype quality (GQ) score of at least 20, at least 10x coverage, a quality by depth (QD) score of at least 2 and a mapping quality (MQ) score of at least 40. Indels were required to have a maximum strand bias (FS) of 200 and a minimum read position rank sum (RPRS) of -20. SNVs were restricted according to VQSR tranche (calculated using the known SNV sites from HapMap v3.3, dbSNP, and the Omni chip array from the 1000 Genomes Project): the cutoff was a tranche of 99.9%. Variants were excluded if marked by EVS as being failures [28]. Variants were annotated to Ensembl 73 using SnpEff [29].

Only genetically European ethnicity samples were included in the analysis. Samples were screened with KING [30] to remove second-degree or higher relatives; samples with incorrect sexes according to X:Y coverage ratios were removed, as were contaminated samples according to VerifyBamID [31].

### Statistical and informatic analysis

Our study used a gene-based collapsing methodology as previously described [6]. For each gene, each sample was indicated as carrying or not carrying a qualifying variant. Qualifying variants were defined for a dominant model requiring at least one qualifying variant per gene with a minor allele frequency (MAF) cutoff of 0.1% internally and 0.01% in each population of the ExAC database, which has been available for longer than the gnomad database[32]. These allele frequency thresholds used a leave-one-out method for the combined sample of cases and controls (where the MAF of each variant was calculated using all samples except for the sample in question). We performed analyses of CCDS genes using two methods to identify qualifying variants: 1) all non-synonymous and canonical splice variants (coding model), and 2) all canonical splice and non-synonymous coding variants except those predicted by PolyPhen-2 HumDiv [33] to be benign (likely gene disrupting model). Qualifying variants were identified using Analysis Tools for Annotated Variants v.6.0 (http://redmine.igm.cumc.columbia.edu/projects/atav/wiki). This is an in-house software package maintained by the Institute for Genomic Medicine (IGM) at Columbia University that compiles genotype and quality information on all cases and controls sequenced at the IGM as well as annotation data from sources like Ensembl, Polyphen, and ExAC in an easy to use format. We also downloaded qualifying variants from the ExAC database [32] using the same criteria for the coding and likely gene disrupting models, restricting to variants that pass ExAC QC and requiring MAFs to be below 0.01% in each population of the ExAC database. While individual-level data are not available from ExAC, making it impossible to determine exactly how many people had a variant in each gene, we used a conservative estimate by adding up the number of carriers of each rare variant in each gene, and we calculated the proportion of people with a qualifying variant in each gene by assuming that no one had more than one variant in the gene. In addition to comparing the numbers of cases and controls with qualifying variants for each model in each gene, we also performed an aggregate collapsing analysis that compared the number of cases and controls with a qualifying variant in any of the known dystonia genes, considered collectively.

Candidate dystonia genes were identified as OMIM genes containing the word “dystonia” in the clinical synopsis plus *FTL (NBIA3), PANK2 (NBIA1), PLA2G6 (NBIA2), ATN1, HTT, PARK2, TAF1 (DYT3), TOR1A, ARSG,* and *CIZ1* (n =148 genes). Isolated dystonia genes were those covered in a recent dystonia review (n =14 genes)[9]. Overall, 94.9% (88.7%) of the coding bases had at least 10x coverage in our control (case) samples, indicating that the sequence data would be able to pick up most causal variants in these genes if they existed. There were 7 genes that had <50% of their bases with at least 10x coverage (<50% coverage in cases and controls: *ARX*, *GJC2,* and *SDHAF1*; poor coverage in cases only: *TSFM, CACNA1B, CACNA1A,* and *PDX1)*. For the aggregate collapsing analysis across all candidate genes, a Fisher’s exact test was performed using http://www.langsrud.com/fisher.htm. Power was calculated using the Genetic Power Calculator [34], with a dystonia prevalence of 1 in 10,000, 80% power, a D-prime of 1, marker allele frequency of 3% (to reflect the frequency of qualifying coding mutations occuring in any of the isolated dystonia genes in the general population) and equal to risk allele frequency, Aa relative risk equal to AA relative risk (dominant model), 20 cases, a control:case ratio of 118.6 (for 2,372 controls), and an alpha of 0.05 for the one test of all isolated dystonia genes in aggregate.

## RESULTS

To characterize the prevalence of individuals with genetic variants that could potentially be described as pathogenic in genes associated with dystonia, we first sequenced the exomes of 2,372 population controls of European ancestry. We additionally sequenced the exomes of 20 patients with sporadic dystonia and of European ancestry, whose clinical characteristics are summarized in Table 1.

**Table 1.**
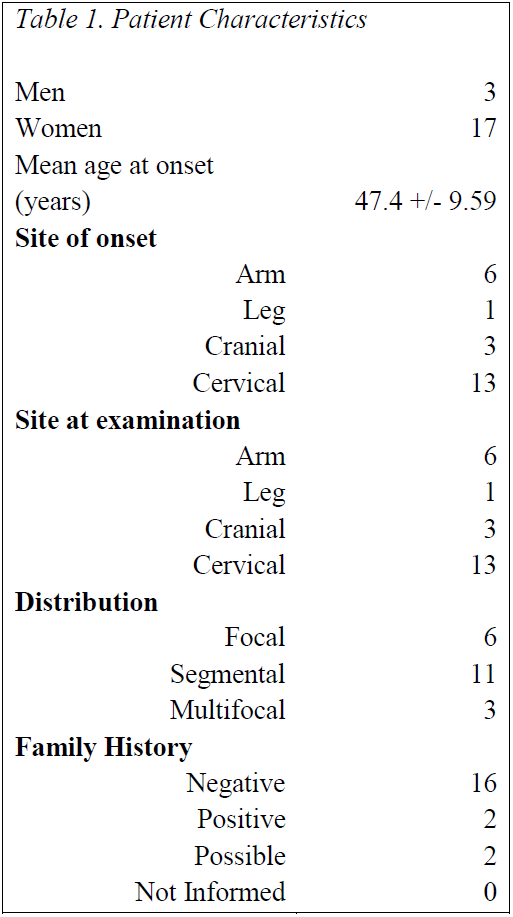
Patient Characteristics.

|  |  |
| --- | --- |
| Men | 3 |
| Women | 17 |
| Mean age at onset<br>(years) | 47.4 +/- 9.59 |
| **Site of onset** |  |
| Arm | 6 |
| Leg | 1 |
| Cranial | 3 |
| Cervical | 13 |
| **Site at examination** |  |
| Arm | 6 |
| Leg | 1 |
| Cranial | 3 |
| Cervical | 13 |
| **Distribution** |  |
| Focal | 6 |
| Segmental | 11 |
| Multifocal | 3 |
| **Family History** |  |
| Negative | 16 |
| Positive | 2 |
| Possible | 2 |
| Not Informed | 0 |

Two sets of “dystonia-associated” genes were curated for the analysis. The first is a more stringent set of genes (14 in total) that have been associated with dystonia in isolation from other phenotypes, “isolated dystonia” (reviewed in [9]). The second set of genes (n=148) more broadly considered those genes in which dystonia had been described as part of the clinical phenotype (see Methods for further details). We first performed a gene-based collapsing analysis where each sample was coded to indicate whether they had a qualifying variant in each sequenced gene. “Qualifying” was defined based on two different genetic models: “coding” and “likely gene disrupting”. We then identified the total number of subjects with a qualifying variant in any of the dystonia genes analysed to arrive at the overall frequency.

Among the group of isolated dystonia genes, we found that 4.2% of controls had qualifying variants in the coding model and 3.0% in the likely gene disrupting model (Table 2). At this rate, we would have expected fewer than 1 of the 20 dystonia cases to have qualifying variants in at least one of these genes by chance, and we found that 0 had such variants. Though we only had 20 cases, our aggregate analysis across all known isolated dystonia genes had 80% power to identify a signal where at least 15% of sporadic dystonia cases were due to pathogenic or possibly pathogenic coding variants in these genes.

**Table 2.** Characteristics of the 2,372 controls used in this study.

|  | n (%) |
| --- | --- |
| **Phenotype** |  |
| Bone disease | 3 (0.1%) |
| Cardiovascular disease | 70 (3%) |
| Dementia caused by APOE 4/4 genotype | 5 (0.2%) |
| Extreme birth weight | 39 (1.6%) |
| Healthy family member of patient with bone, endocrine, hematological, metabolic, immune, or ophthalmic disease | 36 (1.5%) |
| Healthy family member of patient with brain malformation | 259 (10.9%) |
| Healthy family member of patient with epilepsy | 548 (23.1%) |
| Healthy family member of patient with neuropsychiatric disease | 590 (24.9%) |
| Healthy family member of patient with other neurodevelopmental or neurological disease | 99 (4.2%) |
| Healthy volunteer | 124 (5.2%) |
| Hematological disease | 18 (0.8%) |
| Immune disorder | 23 (1%) |
| Infectious disease | 310 (13.1%) |
| Kidney, urological, or endocrine disease | 32 (1.3%) |
| Liver disease | 77 (3.2%) |
| Neurosensory disorder | 3 (0.1%) |
| Ophthalmic disease | 9 (0.4%) |
| Pulmonary disease | 127 (5.4%) |
| Bone disease | 3 (0.1%) |
| **Sex** |  |
| Male | 1279 (53.9%) |
| Female | 1093 (46.1%) |
| **Self-reported ethnicity\*** |  |
| White | 1920 (80.9%) |
| Middle Eastern | 101 (4.3%) |
| Hispanic | 13 (0.5%) |
| Other or unknown | 338 (14.2%) |
| **Sequencing method** |  |
| Whole genome | 375 (15.8%) |
| 37 MB Agilent All Exon | 103 (4.3%) |
| 50 MB Agilent All Exon | 145 (6.1%) |
| 65 MB Agilent All Exon | 648 (27.3%) |
| Nimblegen SeqCap EZ V2.0 or 3.0 Exome Enrichment Kit | 1101 (46.4%) |

\*All controls were genetically confirmed as being of white ethnicity. Ages of controls are not known.

We then broadened our scope to all genes previously reported as associated with dystonia in OMIM and literature reviews as compared to controls, 148 genes in total. We found that 50.6% of controls had qualifying coding variants in at least one of these genes. The frequency among cases was remarkably similar, 45.0%. For the likely gene disrupting model, qualifying variants were found in 37.1% of controls and again, a similar frequency of 35.0% in cases.

Although the primary outcome for this analysis was the aggregated contribution of rare variation in dystonia genes when considered as a group, in Table 3 we also indicate the proportion of cases and controls with qualifying variants for each gene analysed. The dystonia-associated gene that was most often mutated in population controls was *DST*, with 4.1% of controls having variants in the coding model and 1.6% in the likely gene disrupting model. Additionally, all of the primary dystonia genes had qualifying variants in at least one of our controls under the coding model, while 4.7% of the genes in the extended candidate dystonia gene list were never mutated in this control set.

**Table 3.**
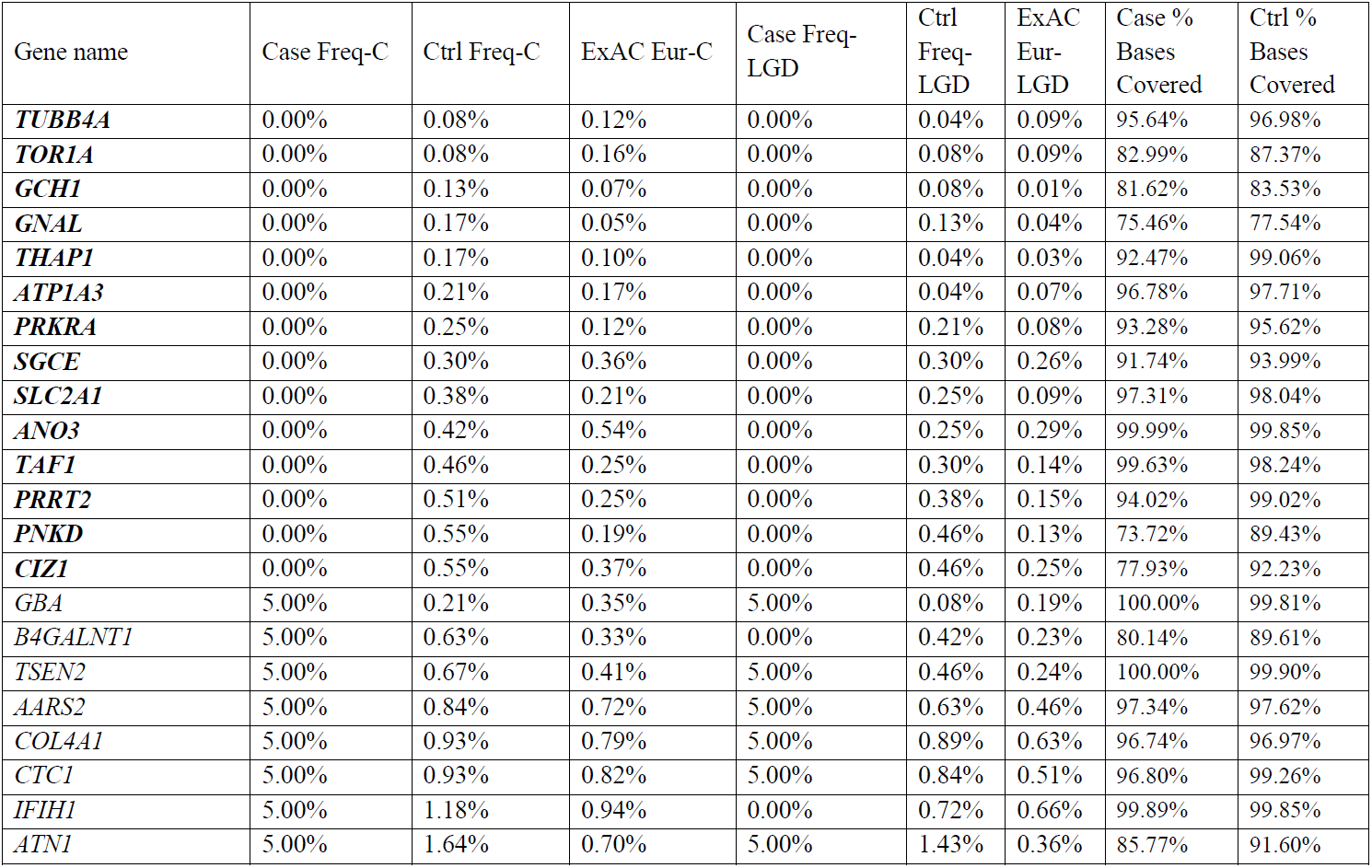
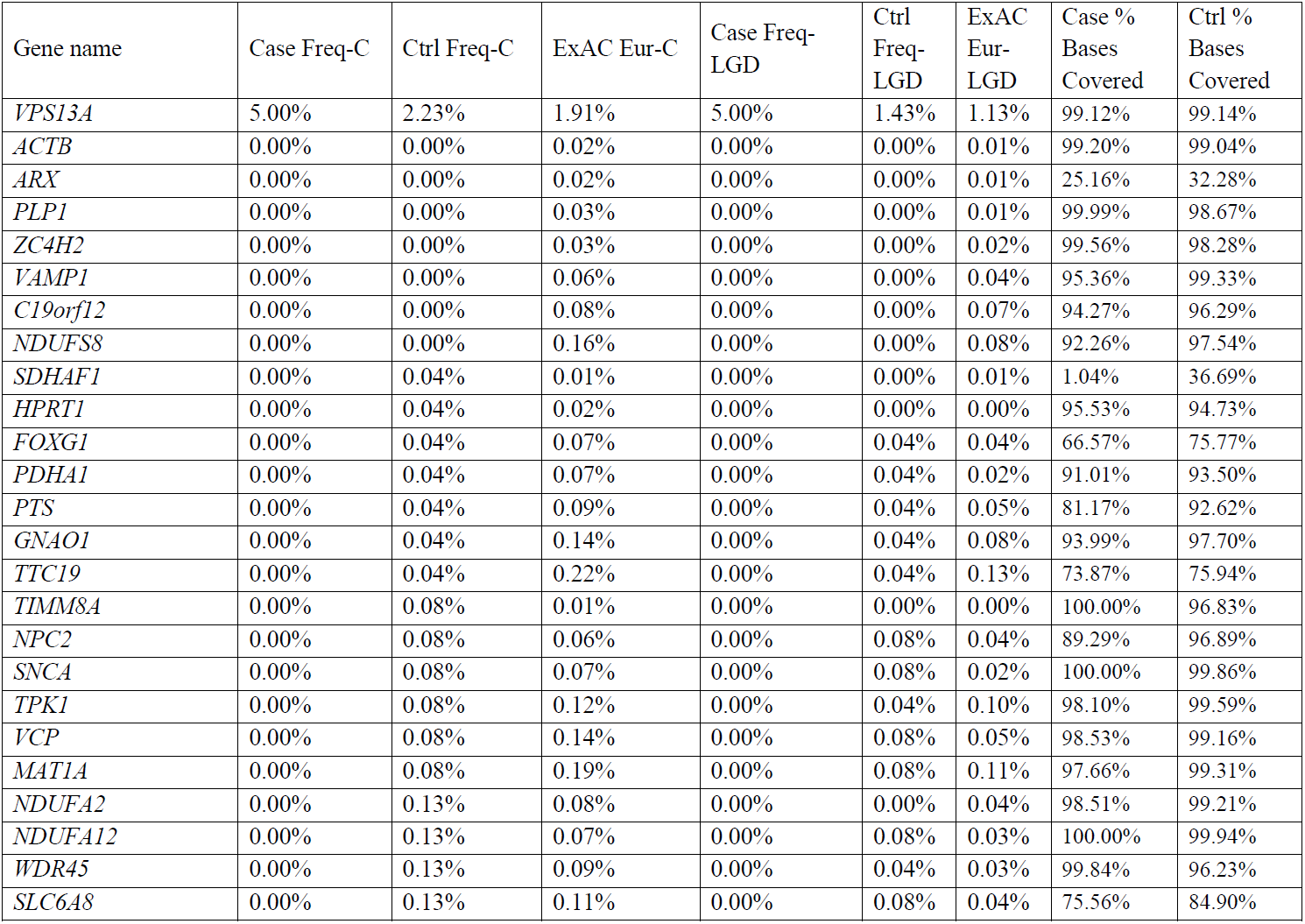
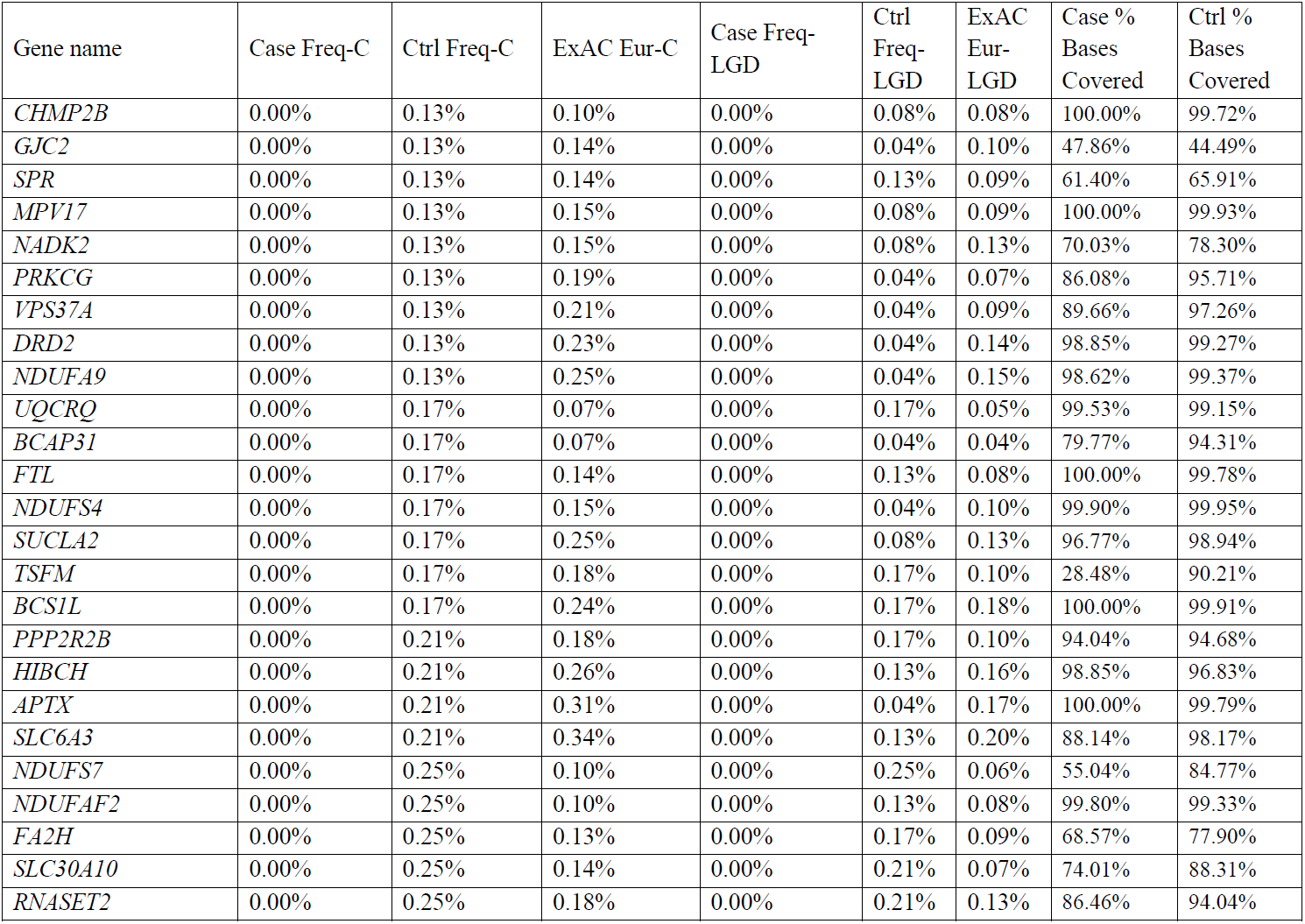
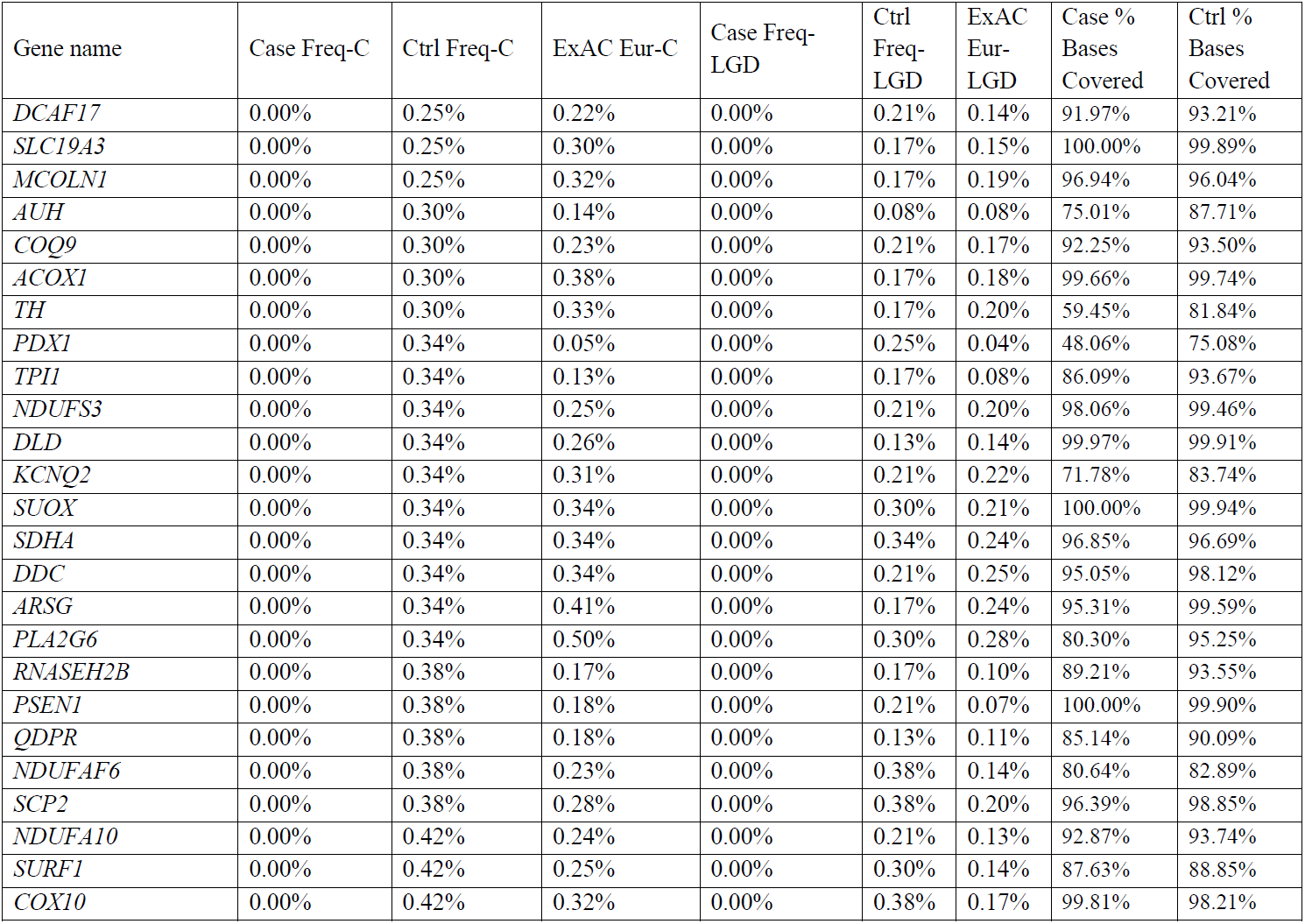
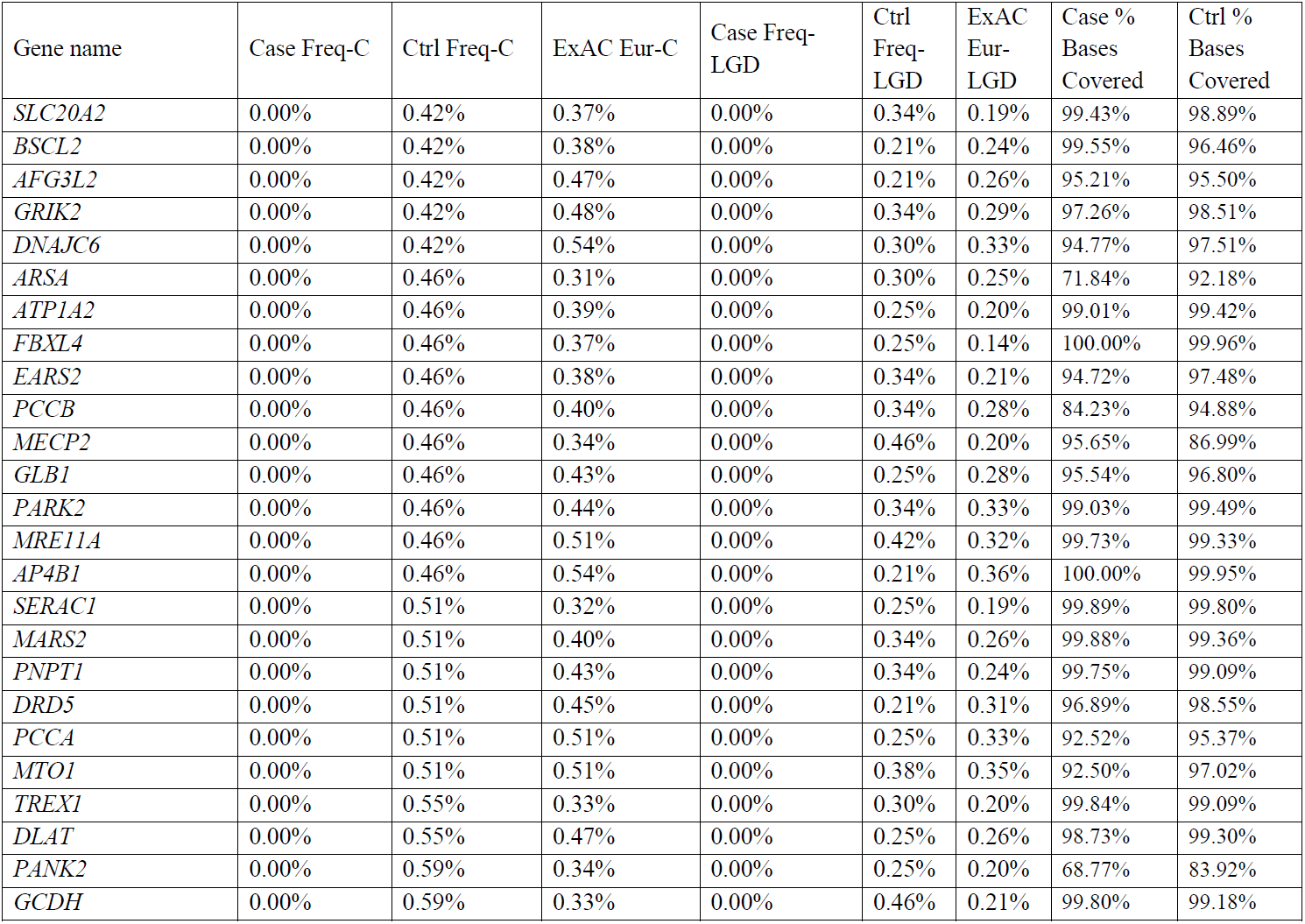
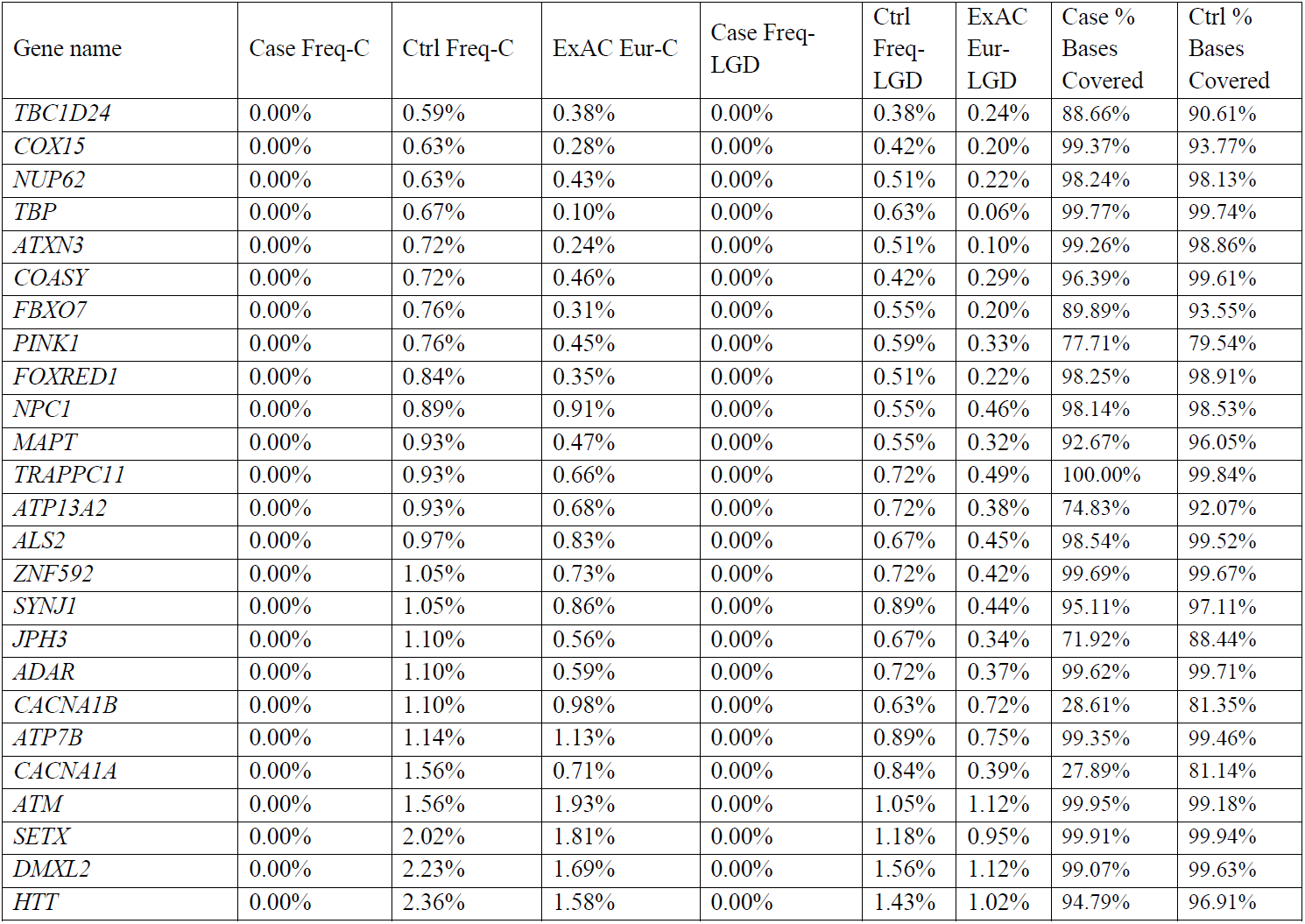
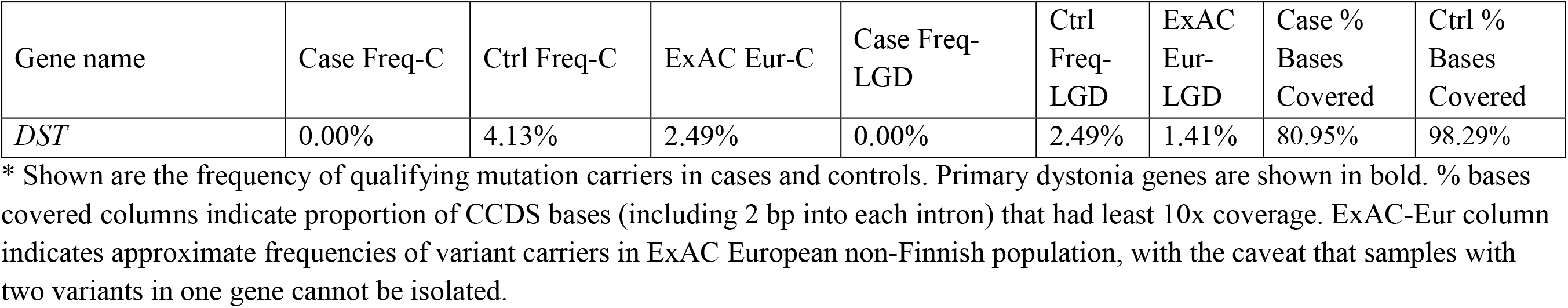
Percent (%) samples with qualifying variants in each of the previously reported dystonia genes for the coding (C) and likely gene disrupting (LGD) models.

Table 3 and S1 Table also present an estimate for the proportion of people in the general population from several ethnicities who have qualifying variants in each of these genes. The data for this section come from the ExAC database, a collection of exome sequence data from >60,000 people at the time of access [32]. Unlike the data presented for our locally sequenced controls, the proportion of carriers of qualifying variants in each ExAC population is an estimate because individual-level data are not available from ExAC; that is, if the same person has two qualifying variants in the same gene, then you will accidentally count them twice. In our locally sequenced controls, we found that 0.5% and 0.9% of controls had more than one qualifying variant in a single candidate dystonia gene under the coding and likely gene disrupting models, respectively. This result indicates that it is uncommon for a person to have multiple qualifying variants in a single candidate dystonia gene, and so the frequency of qualifying variants per gene from ExAC is fairly accurate, which is also supported by the strong similarity between the frequencies of qualifying variants for each gene in our locally sequenced controls and the European ExAC population (S1 Fig and S2 Fig). However, it is important to note that one cannot use ExAC data to estimate how often a person would have a qualifying mutation in any of the genes, i.e., considered in aggregate. In our controls, 15.5% of individuals had rare qualifying variants in more than one dystonia gene (of the 148 dystonia genes) for the coding model, and 8.5% for the likely gene disrupting model.

## DISCUSSION

Here, we have performed exome sequencing and evaluated the frequency of individuals with rare variation in previously reported dystonia genes in 2,372 population controls and a pilot sample of 20 sporadic dystonia patients. We found that it is fairly common for population controls to have likely gene disrupting variants in these genes. Depending on the variant-calling model used, we found that between 37 and 51% of the control population had qualifying variants in dystonia genes. Among the subset of 14 genes associated with causing only dystonia and not being part of a larger syndrome (“isolated”), we found that 3-4% of the population harbored qualifying variants. These frequencies are noteworthy since dystonia is a rare disease, estimated to affect significantly fewer than 0.1% of the population [35]. Thus, the results of our study provide an important point-of-reference for providers in the clinical setting that are considering the significance of a rare variant found in an individual with dystonia.

For the purpose of counseling a patient, knowing that there is not a strong enrichment of variants in dystonia genes in sporadic patients promotes using caution when assigning causality to a variant. Based on our analysis, if all individuals with a qualifying rare variant in a dystonia gene were to be clinically considered for the presence of dystonia, at most 1 individual in 2000 subjects with rare variants in isolated dystonia genes would be expected to have dystonia based on an estimated clinical incidence of 1 in 10,000 and a frequency of such variants in the general population of 3% (and <15% of sporadic dystonia cases). Conversely, for an individual with clinically confirmed dystonia, the chance that their rare variant is incidental is 1 in 25-33 for isolated dystonia genes and 1 in 2-3 for any dystonia-associated gene.

A number of isolated cases of sporadic disease have been reported that possess rare, likely gene-disrupting mutations in previously described dystonia genes [10-13, 15, 17-19], and more recently, the collective burden of rare variation has been assessed in single dystonia genes of interest [11, 19, 21, 36]. The present study is the first to formally describe the overall frequency of rare dystonia gene variants. We focused the study on population controls, but also systematically searched for evidence of enrichment in sporadic dystonia cases. Though our case sample size was small, the collapsing nature of this analysis, in which all dystonia genes are considered as a group, was sufficiently powered to ascertain whether a substantial fraction of sporadic dystonia cases (>15% or 3 subjects) could be explained by rare missense variation in isolated dystonia genes. We found no evidence for an enrichment of rare functional variants in dystonia genes among sporadic dystonia patients. Of course, specific variants and/or individual dystonia-associated genes may still play a role in some cases and larger scale studies are necessary to determine their significance. Our study subjects had predominantly adult-onset focal dystonias, while a recent report indicates a potentially larger contribution of rare variants in dystonia genes to early-onset dystonia [37]. Larger scale studies are necessary to determine these possibilities. Our results indicate however, that this genetic mechanism, on its own, is not sufficient to explain the bulk of sporadic cases and emphasize the need for a better understanding of other potential genetic and non-genetic factors that give rise to dystonia in the population with late-onset sporadic presentation.

We also note that our conclusions specifically apply to rare coding variants and not other sources of genetic variation. For example, exome sequencing does miss many insertions and deletions as well as structural variants and non-coding variants. Causal variants in some of the known dystonia genes, such as the intronic retrotransposon insertion in *TAF1* (DYT3)[38] and some of the larger deletions in SGCE [39, 40], cannot therefore be accurately screened in our study.

Finally, we present the approximate rate at which controls in the ExAC database have rare coding variants in dystonia genes (Table 3 and S1 Table). These numbers are approximate because individual-level data are not available from databases like ExAC and gnomad, and thus proportions will be inflated as you cannot distinguish a situation where one person has two rare functional variants from a situation where two people each have one rare functional variant. This is especially true if one attempts to calculate what proportion of people will have a mutation in any one of a set of genes, in aggregate. For example, adding up all the control carrier frequencies for the 148 genes in Table 3 would indicate that 71.2% of people have qualifying variants in these genes under the coding model, but because 15.5% have mutations in two or more genes, it is actually only 50.5% who have a mutation in at least one of these genes. As a final caveat, all sequencing data sets have their own particular parameters in terms of, for example, the kit used, the sequencing quality, the processing software used, and the quality and annotation cutoffs used to identify variants of interest. It is therefore imprecise to compare aggregate data across multiple variants or genes from datasets such as ExAC or gnomad to one’s own data, and the gold standard will always be to compare sequenced cases to a large number of controls that have been sequenced by the same group using identical methods. It is especially pertinent to be aware of the coverage statistics, as some exons or sites may have poor or no coverage in some datasets.

In conclusion, here we present the first systematic analysis of the aggregated contribution of rare variation in dystonia genes. Our analysis provides a statistical framework that can be used as the field continues to develop more advanced and accurate abilities to ascertain the significance of rare genetic variants in rare diseases. Such ongoing efforts include *in vitro* functional phenotyping and the assembly of genotype-phenotype clinical databases (MDSgene) [41, 42].

Our findings provide an easily accessible resource (Table 3) and highlight the necessity of considering general population frequencies when evaluating the significance of rare variants to rare diseases. The approach, in principle, is also applicable to other rare diseases as genome sequence data increasingly enter the equation of care for an individual patient.

## Acknowledgements

We are grateful to the patients and healthy volunteers who participated in this research. We thank Changrui Xiao for technical assistance and David Goldstein for advice. We would like to acknowledge the following individuals or groups for the contributions of control samples: K. Welsh-Bomer, C. Hulette; W. Lowe; D. Marchuk; S. Schuman, E. Nading; J. Burke; S. Palmer; J. Milner; P. Lugar; C. Moylan; A. M. Diehl; M. Abdelmalek; DUHS (Duke University Health System) Nonalcoholic Fatty Liver Disease Research; M. Winn, R. Gbadegesin; A. Holden; D. Levy; E. Behr; D. Daskalakis; R Buckley; E. Holtzman; M. Hauser; J.Hoover-Fong, N. L. Sobreira and D. Valle; A. Poduri; T. Young and K. Whisenhunt; Z. Farfel, D. Lancet, and E. Pras; G. Cavalleri; N. Delanty; G. Nestadt; J. Samuels, Y. Wang; V. Shashi; M. Carrington; S. Kerns, H. Oster; The Murdock Study Community Registry and Biorepository; C. Woods, K. Schmader, S. MacDonald, M. Yanamadala, H. White, and Crosdaile Retirement Communities; National Institute of Allergy and Infectious Diseases Center for HIV/AIDS Vaccine Immunology (CHAVI) (U19-AI067854), National Institute of Allergy and Infectious Diseases Center for HIV/AIDS Vaccine Immunology and Immunogen Discovery (UM1-AI100645); and the Epi4K Consortium and Epilepsy Phenome/Genome Project.

## Authors’ Roles

E.T.C. (1B,C; 2A,B,C; 3A,B) P.H. (1C; 3B), J.J. (1B,C; 3B), K.T. (1C; 3B), S.T. (1C; 3B), Z.C. (1C; 3B), B.S. (1C, 3B), M.S. (1C, 3B), N.C. (1A,B,C; 2C; 3A,B)

## Financial Disclosures of all authors (for the preceding 12 months)

N.C. receives royalty payment from Circuit Therapeutics (Redwood City, California), serves on the Tourette Syndrome Association scientific advisory board, receives honoraria solely from non-profit academic organizations and has received grants from NINDS, NIMH, Tyler’s Hope for a Dystonia Cure, Bachmann Strauss Dystonia Parkinson Foundation, Dystonia Medical Research Foundation, McKnight Foundation, and the Harrington Discovery Institute. E.T.C. receives support from the National Institute of Mental Health of the National Institutes of Health under award number K01MH098126 and has received grant support from Biogen, Idec. P.H. has consulted for and received grant support from Medtronic, Allergan, Merz, and GE Medical. P.H. serves as Fellowship director of the Duke Movement Disorders Center, which has received funding from Medtronic, Allergan, and Boston Scientific. M.S. reports grant support from the Michael J. Fox Foundation, Parkinson Study Group and has received compensation as a consultant for Eli Lilly, Merz, Osmotica, ProStrakan and SK Life Sciences. He serves on protocol steering committees for Allergan and Biotie, and receives royalties for the “Handbook of Dystonia”. B.S. received grants for clinical trials from Merz, Acadia, Auspex, and CHDI. No other disclosures are reported.

## Supporting information

S1 Fig. Correlation between proportion of samples with samples with qualifying variants in each gene under the coding (C) model in our locally sequenced controls and in the European non-Finnish population in ExAC (r2=0.85; p<0.001).

S2 Fig. Correlation between proportion of samples with samples with qualifying variants in each gene under the likely gene disrupting (LGD) model in our locally sequenced controls and in the European non-Finnish population in ExAC (r2=0.80; p<0.001).

**S1 Table**. Approximate percent (%) ExAC samples with qualifying variants in each of the previously reported dystonia genes for the coding (C) and likely gene disrupting (LGD) models.

## References

1. Do R, Kathiresan S, Abecasis GR. Exome sequencing and complex disease: practical aspects of rare variant association studies. Hum Mol Genet. 2012;21(R1):R1-9. doi:10.1093/hmg/dds387. PubMed PMID: 22983955; PubMed Central PMCID: PMC3459641.

2. Hu H, Roach JC, Coon H, Guthery SL, Voelkerding KV, Margraf RL, et al. A unified test of linkage analysis and rare-variant association for analysis of pedigree sequence data. Nat Biotechnol. 2014;32(7):663-9. doi:10.1038/nbt.2895. PubMed PMID: 24837662; PubMed Central PMCID: PMC4157619.

3. Lee S, Teslovich TM, Boehnke M, Lin X. General framework for meta-analysis of rare variants in sequencing association studies. Am J Hum Genet. 2013;93(1):42-53. doi:10.1016/j.ajhg.2013.05.010. PubMed PMID: 23768515; PubMed Central PMCID: PMC3710762.

4. Cirulli ET, Goldstein DB. Uncovering the roles of rare variants in common disease through whole-genome sequencing. Nat Rev Genet. 2010;11(6):415-25. doi:10.1038/nrg2779. PubMed PMID: 20479773.

5. Vissers LE, de Ligt J, Gilissen C, Janssen I, Steehouwer M, de Vries P, et al. A de novo paradigm for mental retardation. Nat Genet. 2010;42(12):1109-12. doi:10.1038/ng.712. PubMed PMID: 21076407.

6. Cirulli ET, Lasseigne BN, Petrovski S, Sapp PC, Dion PA, Leblond CS, et al. Exome sequencing in amyotrophic lateral sclerosis identifies risk genes and pathways. Science. 2015. doi:10.1126/science.aaa3650. PubMed PMID: 25700176.

7. Do R, Stitziel NO, Won HH, Jorgensen AB, Duga S, Angelica Merlini P, et al. Exome sequencing identifies rare LDLR and APOA5 alleles conferring risk for myocardial infarction. Nature. 2015;518(7537):102-6. doi:10.1038/nature13917. PubMed PMID: 25487149; PubMed Central PMCID: PMC4319990.

8. Albanese A, Bhatia K, Bressman SB, Delong MR, Fahn S, Fung VS, et al. Phenomenology and classification of dystonia: a consensus update. Mov Disord. 2013;28(7):863-73. doi:10.1002/mds.25475. PubMed PMID: 23649720; PubMed Central PMCID: PMC3729880.

9. Lohmann K, Klein C. Genetics of dystonia: what's known? What's new? What's next? Mov Disord. 2013;28(7):899-905. doi:10.1002/mds.25536. PubMed PMID: 23893446.

10. Calakos N, Patel VD, Gottron M, Wang G, Tran-Viet KN, Brewington D, et al. Functional evidence implicating a novel TOR1A mutation in idiopathic, late-onset focal dystonia. Journal of medical genetics. 2010;47(9):646-50. doi:10.1136/jmg.2009.072082. PubMed PMID: 19955557; PubMed Central PMCID: PMC2891583.

11. Dufke C, Sturm M, Schroeder C, Moll S, Ott T, Riess O, et al. Screening of mutations in GNAL in sporadic dystonia patients. Mov Disord. 2014;29(9):1193-6. doi:10.1002/mds.25794. PubMed PMID: 24408567.

12. Groen JL, Andrade A, Ritz K, Jalalzadeh H, Haagmans M, Bradley TE, et al. CACNA1B mutation is linked to unique myoclonus-dystonia syndrome. Hum Mol Genet. 2015;24(4):987-93. doi:10.1093/hmg/ddu513. PubMed PMID: 25296916.

13. Hettich J, Ryan SD, de-Souza ON, Saraiva Macedo Timmers LF, Tsai S, Atai NA, et al. Biochemical and cellular analysis of human variants of the DYT1 dystonia protein, 18 TorsinA/TOR1A. Hum Mutat. 2014;35(9):1101-13. doi:10.1002/humu.22602. PubMed PMID: 24930953; PubMed Central PMCID: PMC4134760.

14. Kock N, Naismith TV, Boston HE, Ozelius LJ, Corey DP, Breakefield XO, et al. Effects of genetic variations in the dystonia protein torsinA: identification of polymorphism at residue 216 as protein modifier. Human molecular genetics. 2006;15(8):1355-64. doi:10.1093/hmg/ddl055. PubMed PMID: 16537570.

15. Lohmann K, Uflacker N, Erogullari A, Lohnau T, Winkler S, Dendorfer A, et al. Identification and functional analysis of novel THAP1 mutations. Eur J Hum Genet. 2012;20(2):171-5. doi:10.1038/ejhg.2011.159. PubMed PMID: 21847143; PubMed Central PMCID: PMC3260936.

16. Mencacci NE, R'Bibo L, Bandres-Ciga S, Carecchio M, Zorzi G, Nardocci N, et al. The CACNA1B R1389H variant is not associated with myoclonus-dystonia in a large European multicentric cohort. Hum Mol Genet. 2015. doi:10.1093/hmg/ddv255. PubMed PMID: 26157024.

17. Nibbeling E, Schaake S, Tijssen MA, Weissbach A, Groen JL, Altenmuller E, et al. Accumulation of rare variants in the arylsulfatase G (ARSG) gene in task-specific dystonia. J Neurol. 2015;262(5):1340-3. doi:10.1007/s00415-015-7718-3. PubMed PMID: 25825126.

18. Vemula SR, Xiao J, Zhao Y, Bastian RW, Perlmutter JS, Racette BA, et al. A rare sequence variant in intron 1 of THAP1 is associated with primary dystonia. Mol Genet Genomic Med. 2014;2(3):261-72. doi:10.1002/mgg3.67. PubMed PMID: 24936516; PubMed Central PMCID: PMC4049367.

19. Zech M, Gross N, Jochim A, Castrop F, Kaffe M, Dresel C, et al. Rare sequence variants in ANO3 and GNAL in a primary torsion dystonia series and controls. Mov Disord. 2014;29(1):143-7. doi:10.1002/mds.25715. PubMed PMID: 24151159.

20. Takeshita E, Saito Y, Nakagawa E, Komaki H, Sugai K, Sasaki M, et al. Late-onset mental deterioration and fluctuating dystonia in a female patient with a truncating MECP2 mutation. Journal of the neurological sciences. 2011;308(1-2):168-72. doi:10.1016/j.jns.2011.06.008. PubMed PMID: 21722922.

21. Groen JL, Ritz K, Warner TT, Baas F, Tijssen MA. DRD1 rare variants associated with tardive-like dystonia: a pilot pathway sequencing study in dystonia. Parkinsonism & related disorders. 2014;20(7):782-5. doi:10.1016/j.parkreldis.2014.04.002. PubMed PMID: 24768614.

22. Zech M, Jochim A, Boesch S, Weber S, Meindl T, Peters A, et al. Systematic TOR1A nonc.907_909delGAG variant analysis in isolated dystonia and controls. Parkinsonism & related disorders. 2016;31:119-23. doi:10.1016/j.parkreldis.2016.07.013. PubMed PMID: 27477622.

23. Long Y, Chen Y, Qian Y, Wang J, Luo L, Huang X, et al. A rare variant in TOR1A exon 5 associated with isolated dystonia in southwestern Chinese. Mov Disord. 2017;32(7):1083-7. doi:10.1002/mds.27016. PubMed PMID: 28432771.

24. Basu S, Pan W. Comparison of statistical tests for disease association with rare variants. Genetic epidemiology. 2011;35(7):606-19. doi:10.1002/gepi.20609. PubMed PMID: 21769936; PubMed Central PMCID: PMC3197766.

25. Cirulli ET. The Increasing Importance of Gene-Based Analyses. PLoS genetics. 2016;12(4):e1005852. doi:10.1371/journal.pgen.1005852. PubMed PMID: 27055023; PubMed Central PMCID: PMCPMC4824358.

26. Li H, Durbin R. Fast and accurate short read alignment with Burrows-Wheeler transform. Bioinformatics (Oxford, England). 2009;25(14):1754-60. doi:10.1093/bioinformatics/btp324. PubMed PMID: 19451168; PubMed Central PMCID: PMC2705234.

27. McKenna A, Hanna M, Banks E, Sivachenko A, Cibulskis K, Kernytsky A, et al. The Genome Analysis Toolkit: a MapReduce framework for analyzing next-generation DNA sequencing data. Genome Res. 2010;20(9):1297-303. doi:10.1101/gr.107524.110. PubMed PMID: 20644199; PubMed Central PMCID: PMC2928508.

28. ExomeVariantServer. NHLBI GO Exome Sequencing Project (ESP) NHLBI GO Exome Sequencing Project (ESP): Seattle, WA; URL: http://evs.gs.washington.edu/EVS/. Available from: URL: http://evs.gs.washington.edu/EVS/.

29. Cingolani P, Platts A, Wang le L, Coon M, Nguyen T, Wang L, et al. A program for annotating and predicting the effects of single nucleotide polymorphisms, SnpEff: SNPs in the genome of Drosophila melanogaster strain w1118; iso-2; iso-3. Fly (Austin). 2012;6(2):80-92. doi:10.4161/fly.19695. PubMed PMID: 22728672; PubMed Central PMCID: PMC3679285.

30. Manichaikul A, Mychaleckyj JC, Rich SS, Daly K, Sale M, Chen WM. Robust relationship inference in genome-wide association studies. Bioinformatics. 2010;26(22):2867-73. doi:10.1093/bioinformatics/btq559. PubMed PMID: 20926424; PubMed Central PMCID: PMC3025716

31. Jun G, Flickinger M, Hetrick KN, Romm JM, Doheny KF, Abecasis GR, et al. Detecting and estimating contamination of human DNA samples in sequencing and array-based genotype data. American journal of human genetics. 2012;91(5):839-48. doi:10.1016/j.ajhg.2012.09.004. PubMed PMID: 23103226; PubMed Central PMCID: PMC3487130.

32. Lek M, Karczewski KJ, Minikel EV, Samocha KE, Banks E, Fennell T, et al. Analysis of protein-coding genetic variation in 60,706 humans. Nature. 2016;536(7616):285-91. doi:10.1038/nature19057. PubMed PMID: 27535533; PubMed Central PMCID: PMCPMC5018207.

33. Adzhubei I, Jordan DM, Sunyaev SR. Predicting functional effect of human missense mutations using PolyPhen-2. Curr Protoc Hum Genet. 2013;Chapter 7:Unit7 20. doi:10.1002/0471142905.hg0720s76. PubMed PMID: 23315928.

34. Purcell S, Cherny SS, Sham PC. Genetic Power Calculator: design of linkage and association genetic mapping studies of complex traits. Bioinformatics (Oxford, England). 2003;19(1):149-50. PubMed PMID: 12499305.

35. Defazio G. The epidemiology of primary dystonia: current evidence and perspectives. Eur J Neurol. 2010;17 Suppl 1:9-14. doi:10.1111/j.1468-1331.2010.03053.x. PubMed PMID: 20590802.

36. Zech M, Jech R, Havrankova P, Fecikova A, Berutti R, Urgosik D, et al. KMT2B rare missense variants in generalized dystonia. Mov Disord. 2017;32(7):1087-91. doi:10.1002/mds.27026. PubMed PMID: 28520167.

37. Zech M, Boesch S, Jochim A, Weber S, Meindl T, Schormair B, et al. Clinical exome sequencing in early-onset generalized dystonia and large-scale resequencing follow-up. Mov Disord. 2016. doi:10.1002/mds.26808. PubMed PMID: 27666935.

38. Makino S, Kaji R, Ando S, Tomizawa M, Yasuno K, Goto S, et al. Reduced neuron-specific expression of the TAF1 gene is associated with X-linked dystonia-parkinsonism. Am J Hum Genet. 2007;80(3):393-406. doi:10.1086/512129. PubMed PMID: 17273961; PubMed Central PMCID: PMCPMC1821114.

39. Asmus F, Salih F, Hjermind LE, Ostergaard K, Munz M, Kuhn AA, et al. Myoclonus-dystonia due to genomic deletions in the epsilon-sarcoglycan gene. Ann Neurol. 2005;58(5):792-7. doi:10.1002/ana.20661. PubMed PMID: 16240355.

40. Zimprich A, Grabowski M, Asmus F, Naumann M, Berg D, Bertram M, et al. Mutations in the gene encoding epsilon-sarcoglycan cause myoclonus-dystonia syndrome. Nat Genet. 2001;29(1):66-9. doi:10.1038/ng709. PubMed PMID: 11528394.

41. Lill CM, Mashychev A, Hartmann C, Lohmann K, Marras C, Lang AE, et al. Launching the movement disorders society genetic mutation database (MDSGene). Mov Disord. 2016;31(5):607-9. doi:10.1002/mds.26651. PubMed PMID: 27156390.

42. Capetian P, Pauly MG, Azmitia LM, Klein C. Striatal cholinergic interneurons in isolated generalized dystonia-rationale and perspectives for stem cell-derived cellular models. Front Cell Neurosci. 2014;8:|p205. doi:10.3389/fncel.2014.00205. PubMed PMID: 25120431; PubMed Central PMCID: PMCPMC4112996.

